# Chemogenetic approaches reveal dual functions of microglia in epilepsy

**DOI:** 10.1101/2023.05.28.542660

**Authors:** Aastha Dheer, Dale B. Bosco, Jiaying Zheng, Lingxiao Wang, Shunyi Zhao, Koichiro Haruwaka, Min-Hee Yi, Abhijeet Barath, Dai-Shi Tian, Long-Jun Wu

## Abstract

Microglia are key players in maintaining brain homeostasis and exhibit phenotypic alterations in response to epileptic stimuli. However, it is still relatively unknown if these alterations are pro- or anti-epileptic. To unravel this dilemma, we employed chemogenetic manipulation of microglia via of the artificial Gi-Dreadd receptor within a kainic acid (KA) induced murine seizure model. Our results indicate that Gi-Dreadd activation can reduce seizure severity. Additionally, we observed increased interaction between microglia and neuronal soma, which correlated with reduced neuronal hyperactivity. Interestingly, prolonged activation of microglial Gi-Dreadds by repeated doses over 3 days, arrested microglia in a less active, homeostatic-like state, which associated with increased neuronal loss after KA induced seizures. RNAseq analysis revealed that prolonged activation of Gi-Dreadd interferes with interferon β signaling and microglia proliferation. Thus, our findings highlight the importance of microglial activation not only during *status epilepticus* (SE) but also within later seizure induced pathology.

## Introduction

Epilepsy is a brain disorder characterized by recurrent seizures which affects over 50 million people worldwide [1]. Despite the availability of various anti-epileptic drugs (AEDs), one-third epilepsy patients remain refractory to treatment. Since the most of commonly used AEDs preferentially target neuronal mechanisms, it is critical to explore alternative mechanisms to provide better therapeutic strategies [2–4]. Microglia, the immune cells of central nervous system (CNS), are key regulators of brain disease [5, 6]. Microglia can also sense alterations in neuronal signaling via neurotransmitters and neuromodulators and can regulate neuronal functions [7–11].

Accumulating evidence suggests microglia have a significant role in epileptogenesis [3, 6, 12, 13] and neuronal death after *status epilepticus* [13–17]. However, whether microglia are protective or neurotoxic in epilepsy is still under debate and likely dependent on spatial and temporal factors. Nevertheless, the protective roles of microglia are being increasingly recognized. For instance, interactions between microglia and neurons have been shown to reduce neuronal firing and seizure severity [3, 18–22]. The deletion of the P2Y12 receptor, which is critical for ATP sensing, was shown to aggravate seizures [19, 21]. Disruption of microglial fractalkine signaling was also shown to exacerbate seizures [23]. Moreover, depletion of microglia was shown to aggravate seizures both acutely and the development of spontaneous recurrent seizures [13]. This contrasts with other reports suggesting that depleting microglia reduced deleterious effects [24, 25]. Thus, there is need to fully address this interesting duality of microglia function in epilepsy by precisely targeting microglia within a seizure context.

Novel approaches like chemogenetic manipulations using artificial Designer Receptors Exclusively Activated by Designer Drugs (Dreadd) which enables fine modulation of targeted signaling pathways [26, 27]. The most commonly used Dreadds are the human M3 muscarinic receptor (hM3Dq) which enhances activity via Gq signaling [28] and hM4Di which can silence activity via Gi signaling [27]. The most widely used ligand for these receptors is Clozapine-N-Oxide (CNO), a pharmacologically inert metabolite of Clozapine and an atypical antipsychotic drug (Armbruster et al., 2007; Roth et al., 1994). However, several other compounds like compound 21 [29], perlapine [30], and descholoro-clozapine [31] have been designed for increased Dreadd efficacy and specificity.

Previous studies have utilized Dreadd approaches primarily to modulate neuronal activity [32–35]. However, some have utilized chemogenetic approaches within microglia to modulate peripheral-nerve-injury-induced neuropathic pain [36, 37]. Further, recent findings found that microglial Gi-signaling can regulate microglia-neuron interactions and neuronal excitability [38]. Microglia interact with neurons at specialized areas to engage in cell-cell communication [20]. The microglial specific Gi-coupled receptor P2Y12 may have a crucial role in this communication [20, 21]. In the present study we aim to explore the role of microglia in epileptogenesis by specifically manipulating microglial Gi-signaling using a chemogenetic approach.

## Results

### Microglial Gi-Dreadd activation reduces seizure severity in the ICV-KA model of epilepsy

Previous studies have highlighted the protective role of microglia in acute seizure models [3, 13, 21]. However, the mechanism underlying microglial neuroprotection during seizure onset remains largely unknown. To address this question, we used Cx3cr1^CreER/WT^: R26^LSL-hM4Di/WT^ mice (Gi-Dreadd) which specifically express Gi-Dreadd in microglia under the inducible CX3CR1 promoter. Cx3cr1^CreER/WT^ mice (Control) without Gi-Dreadd expression were used as Controls (**Fig. 1A**). **Fig. 1B** illustrates the experimental timeline. Our results revealed that activation of microglial Gi-Dreadd by CNO significantly reduced seizure score (*p* < 0.05) when compared to controls (**Fig. 1C, D**). To validate Gi-Dreadd expression, we performed immunostaining for HA (Dreadd tag) in Gi-Dreadd and Control groups. (**Fig. 1E**). In the absence of tamoxifen injection, no HA expression was observed in Gi-Dreadd mice confirming cre dependent Gi-Dreadd expression (**Suppl. Fig. 1A**). The results showed the expression of HA positive cells (*p* < 0.0001) was specific to Gi-Dreadd mice (**Fig. 1F**). Additionally, to observe Gi-Dreadd cellular specificity, sections were co-stained with GFAP, NeuN, and IBA1 markers for astrocyte, neurons, and microglia respectively (**Fig. 1G**). Indeed, HA expression colocalized with IBA1 staining (*p* < 0.0001) but not with NeuN or GFAP (**Fig. 1H**).

**Figure 1:**
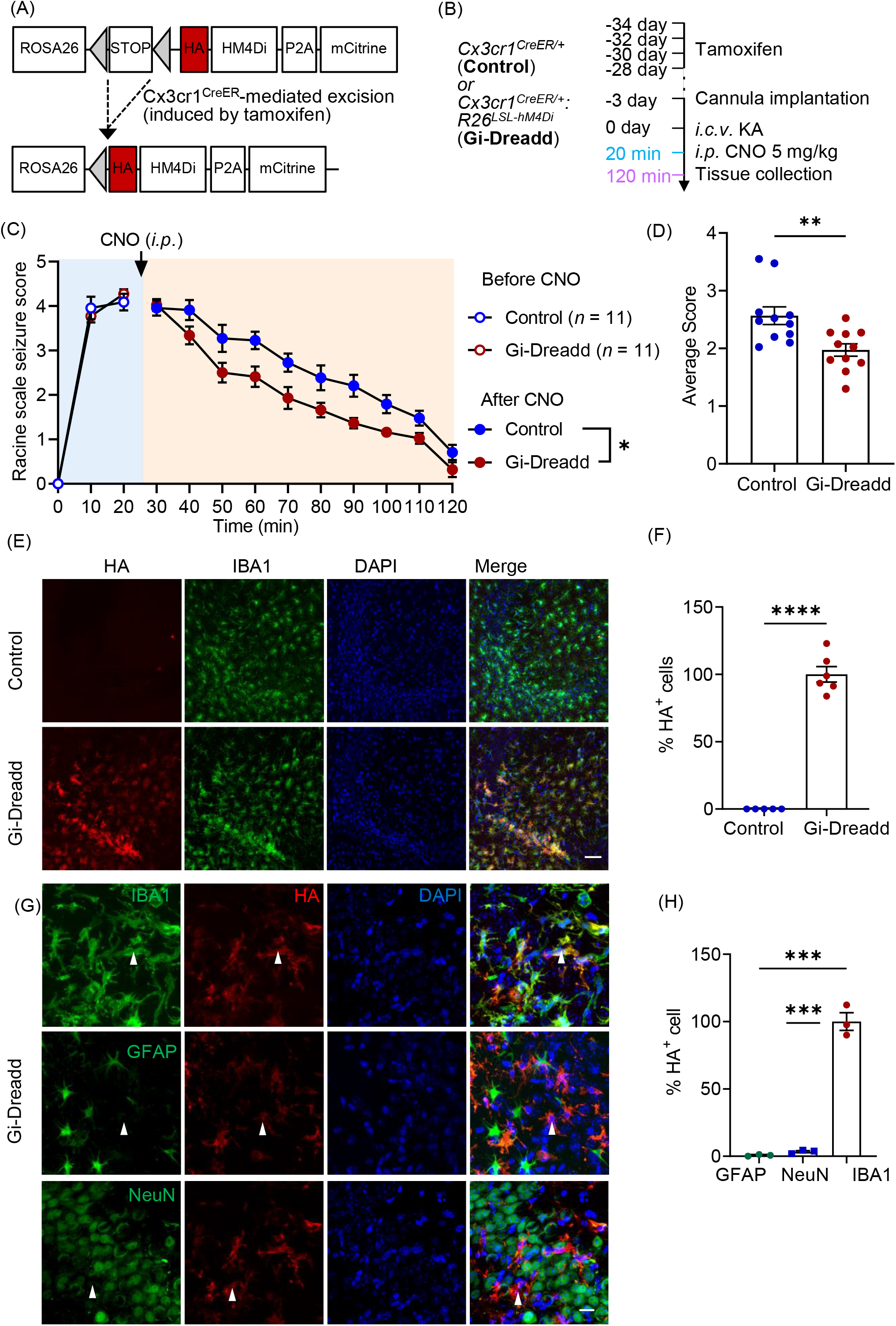
Chemogenetic modulation of microglia reduces seizure severity in the ICV-KA model of epilepsy. (A) Generation of Cx3cr1^creER/WT^: R26^LSL-hM4Di/WT^ transgenic mice. (B) Experimental timeline for i.c.v. KA and CNO (i.p.) injection. (C) Seizure profile for mice that obtained a score of 3 and above in the first 20 min after KA administration. Results indicate that Gi-Dreadd activation by CNO reduced the severity of seizures. The effects were most significant, 30min after CNO administration (n =11/group). Analyzed by two-way ANOVA. (D) Gi-Dreadd significantly reduced average seizure score. Analyzed by t-test. (E) Representative images of IBA1 (green), DAPI (blue), and HA immunostaining (red) in the CA3 region of the hippocampus of Cx3cr1^creER/WT^: R26^LSL-hM4Di/WT^ mice (Gi-Dreadd) after tamoxifen. Scale bar, 50 µm. (F) HA expression co-localized with IBA1+ cells in Gi-Dreadd mice, but not in Cx3cr1^creER/WT^ (control). (G) Representative images of HA tag (red) colocalization with IBA1, NeuN, or GFAP (green) in the hippocampus of Gi-Dreadd mice. Scale bar, 15 µm. (H) Summarized results show HA was only co-localized with IBA1 positive cells. Data represented as mean ± SEM, *p < 0.05, ** p < 0.01, *** p < 0.001.

Finally, since Cx3cr1 is also expressed by peripheral macrophages, we wanted to confirm the observed phenotype was specific to microglial Gi-Dreadd activation. As such, we used the Tmem119^creER^ mouse line to express Gi-Dreadd specifically in microglia [39, 40]. Consistently, we observed a significant reduction in seizure score after CNO injection in the Gi-Dreadd mice when compared with controls (**Suppl. Fig. 1 B, C**). Our results clearly demonstrate that activation of Gi-Dreadd within microglia affects seizures acutely.

### Acute Gi-Dreadd activation increases microglia-neuronal interaction and reduces neuronal hyperactivity following seizures

We next wanted to investigate how Gi-Dreadd activation was affecting microglial behavior. Immunostaining results (**Fig. 2A**) revealed that Gi-Dreadd activation resulted in increased (*p* < 0.05) IBA1^+^ cell number (**Fig. 2B**), and immunoreactivity (*p <* 0.01) (**Fig. 2C**) within the CA3 region 2 h post-KA. Further, Sholl analysis revealed a significant reduction (*p <* 0.0001) overall microglial process complexity within the Gi-Dreadd group when compared to control (**Fig. 2D, E**). Soma size was also significantly increased (*p <* 0.05) in Gi-Dreadd microglia (**Fig. 2F**). In the CA1 region, we observed an increase in IBA1^+^ area (*p <* 0.05) (**Suppl. Fig. 2 A, C**), but there was no difference in microglial cell number (**Suppl. Fig. 2B**) To further characterize microglial alterations, we performed CD68 immunostaining of the CA3 region. The results showed significant (*p <* 0.05) upregulation of CD68 expression in the Gi-Dreadd group when compared to control (**Fig. 2G, H**). Together, these results clearly demonstrate that Gi-Dreadd signaling influences microglial activation in response to KA-induced seizures.

**Figure 2:**
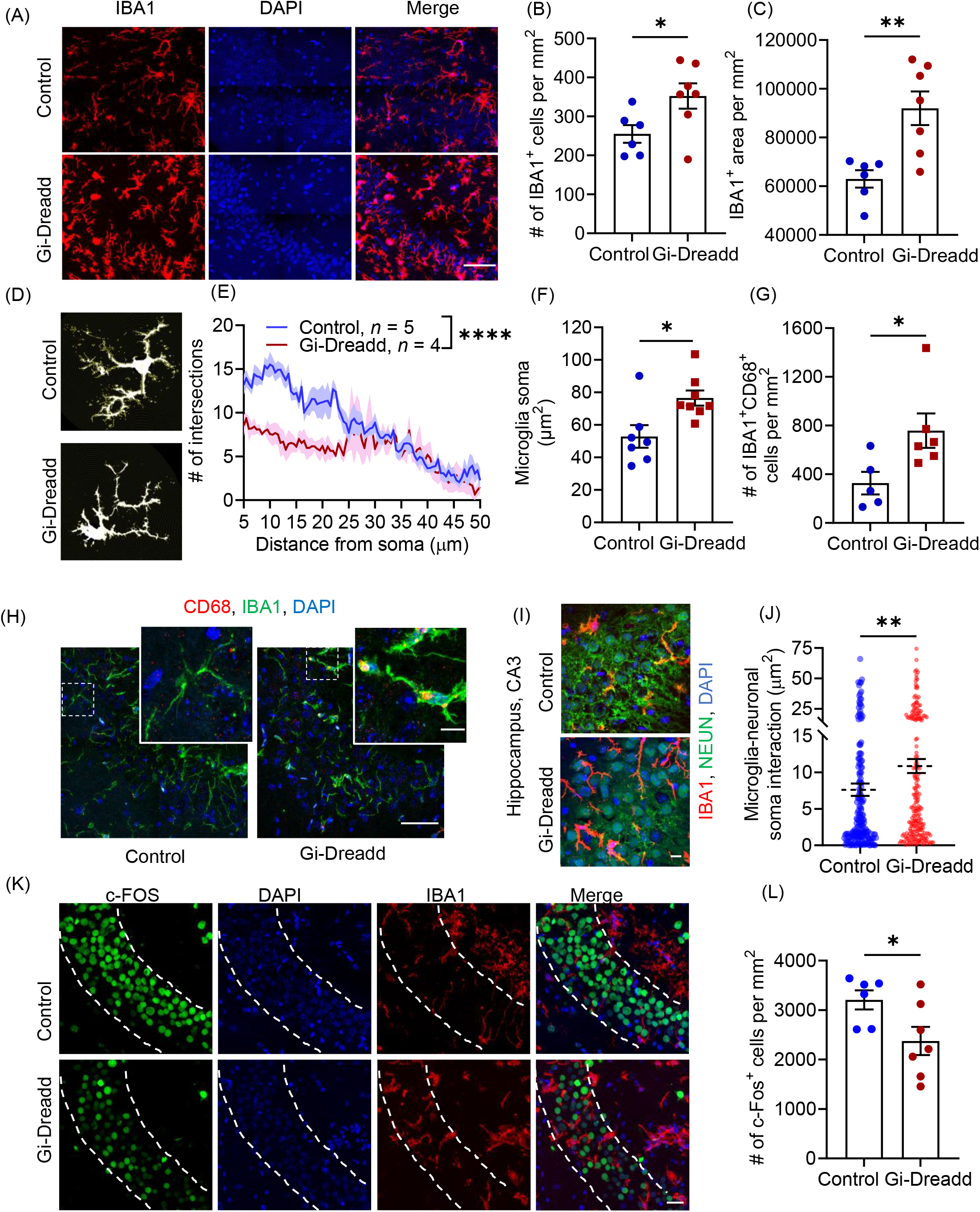
Acute activation of microglial Gi-Dreadd by CNO reduced neuronal hyperactivity following status epilepticus. (A) Representative immunostaining images of IBA1 (red) and DAPI (blue) in CA3 region of hippocampus of Gi-Dreadd and control mice 2 h after KA administration. Scale bar, 50 µm. IBA1 positive cells (B) and IBA1 immunoreactive area (C) were increased in Gi-Dreadd when compared to control. (D) Representative images depicting morphological alterations in microglia. (E) Process complexity as determined by Sholl analysis. Analyzed by two-way ANOVA. (F) Gi-Dreadd activation increased microglial soma size. (G) Gi-Dreadd activation increased number of IBA1+CD68+ cells with respect to controls. (H) Representative images of IBA1 (green), DAPI (blue), and CD68 (red) co-labelling in CA3 region of hippocampus, scale bar, 50 µm. Magnified inset images showing co-localized CD68 expression with IBA1 (yellow), scale bar, 10 µm. (I) Representative images of microglia-neuronal soma interactions, scale bar, 10 µm. (J) Gi-Dreadd activation increased microglia interaction (IBA1+ area) with Neuronal soma (each dot represents individual NeuN+ ROI) when comparison with controls following acute seizures. (K) Representative images of c-FOS (green), IBA1 (red), and DAPI (blue), scale bar, 30 µm. (L) Gi-Dreadd activation reduced c-FOS+ cell number following acute seizures when compared to control. Data represented as mean ± SEM, *p < 0.05, ** p < 0.01, *** p < 0.001, **** p < 0.0001.

To further investigate how microglial Gi-Dreadd activation reduces seizure severity, we examined the interaction between microglia and neurons. Our previous studies found that neuronal hyperactivity after seizures induces microglial process extension towards neuron in the hippocampus [21]. Interestingly, we observed a significant increase (*p <* 0.05) in IBA1^+^ area overlap with NeuN^+^ somas in Gi-Dreadd mice after *status epilepticus* when compared to controls (**Fig. 2 I, J**). To investigate neuronal activity in relation to microglial Gi-Dreadd activation, we analyzed the number of hyperactive c-FOS^+^ neurons present following KA-induced seizures. The results showed a significant decrease (*p <* 0.05) in c-FOS^+^ neurons in the CA3 region of Gi-Dreadd mice when compared to controls (**Fig. 2K, L**). When investigating the CA1 region we did not observe differences in both microglia-neuron interaction after microglial Gi-Dreadd activation, cFOS^+^ neuron number (**Suppl. Fig. 2D-F**). Together, these results indicate that microglial Gi-signaling has a role in regulating microglia-neuron interaction and may be neuroprotective.

### Prolonged Gi-Dreadd activation induction reduces microglia activation and restores homeostatic state

We next wanted to determine how microglial Gi-Dreadd activation affected post-seizure pathology. As such, we examined collected tissues 3 days post-KA when microglial numbers have been reported to peak [4]. To maintain Gi-Dreadd activation, CNO was injected once daily for three days following KA administration (**Fig. 3A**). We carried out immunostaining for IBA1 and CD68 (**Fig. 3B**) in the CA3 region of the hippocampus. The results revealed that upon 3 days after seizures, the number of microglia (**Fig. 3C**) in Gi-Dreadd group was decreased as compared to control group. Additionally, the soma size for Gi-Dreadd microglia was also reduced as compared to the control microglia (**Fig. 3D**). Furthermore, the surface area/volume ratio of microglia is higher in Gi-Dreadd group suggesting that they were more ramified as compared to control group (**Fig. 3E**). The expression of CD68 was also reduced in the Gi-Dreadd group (**Fig. 3F**), indicating decreased microglial reactivity. These results suggest that microglial Gi-Dreadds activation reduces microglial reactivity after 3 days of KA-induced seizures.

**Figure 3:**
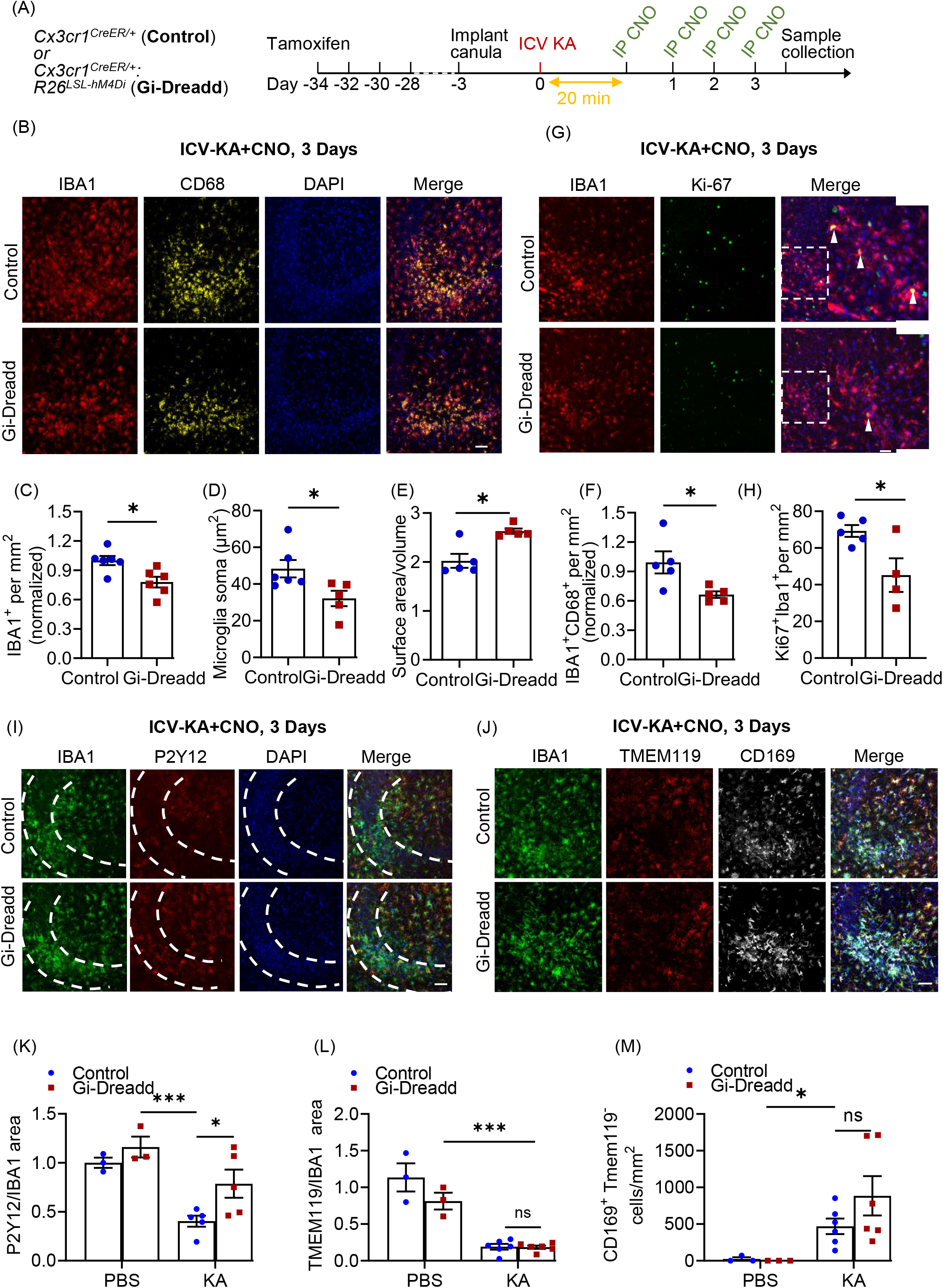
Prolonged activation of Gi-signaling by repeated CNO injections, arrests Gi-Dreadd expressing microglia in homeostatic-like state 3 days after i.c.v. KA induced SE. (A) Experimental timeline. (B) Representative images of IBA1 (red), CD68 (yellow) and DAPI (blue) in the hippocampal CA3 region of Gi-Dreadd and Control mice. Scale bar, 50 µm. Prolonged activation of Gi-Dreadd signalling (C) decreased the number of IBA1+ cells and (D) reduced microglial soma size when compared to controls. (E) Prolonged Gi-Dreadd activation increased morphological complexity of microglia. (F) Prolonged Gi-Dreadd activation reduced CD68 expression. (G) Representative images of IBA1 (red), Ki-67 (green) and DAPI (blue). Scale bar, 50 µm. (H) Prolonged Gi-Dreadd activation reduced the number of Ki-67+IBA1+ cells when compared to controls. (I) Representative images of IBA1 (green), P2Y12 (red) and DAPI (blue), scale bar, 50 µm. (J) Representative images showing infiltrating monocytes in the CA3 region of hippocampus as determined by CD169 (grey), TMEM119 (red), IBA1 (green) and DAPI (blue) staining, scale bar, 50 µm. Cells which express CD169 but lack TMEM119 expression were considered infiltrating macrophages. (K) Prolonged Gi-Dreadd activation increased the ratio of P2Y12 area per IBA1 area when compared to control. (L) No difference in the ratio of Tmem119 area per IBA1 area between groups. (M) No difference in the number of CD169+TMEM119-cells between groups. Data represented as mean ± SEM, *p < 0.05, ** p < 0.01, *** p < 0.001.

We also looked for microglia proliferation following 3 days of i.c.v.-KA and CNO injections using Ki-67 (marker for proliferating cells) co-labelled with IBA1 (**Fig. 3G**). The results showed a significant decrease (*p* < 0.05) in no. of proliferating microglial cells in the Gi-Dreadd group as compared to the controls (**Fig. 3H**).

Next, we assessed the expression of P2Y12, a microglial homeostatic marker. Our results revealed a significant downregulation (*p <* 0.001) of P2Y12 following seizure induction when compared to sham controls (**Fig. 3I, K, suppl. 3A**). Interestingly, P2Y12 down-regulation was significantly suppressed by Gi-Dreadd activation (*p <* 0.05) (**Fig. 3I, K**). Additionally, we investigated expression of TMEM119, another microglial homeostatic marker. However, our results revealed that while seizure induction reduced TMEM119 expression, there was not a statistically significant differences between Gi-Dreadd and control groups (**Fig. 3J, L**).

Finally, we wanted to investigate how microglial Gi Dreadd activation affected monocyte infiltration. It is well established that following *status epilepticus* there is significant infiltration of peripheral monocytes that can have important roles in epilepsy associated neurodegeneration [4, 17, 41]. As such, we combined TMEM119 and CD169 immunostaining to differentiate microglia from infiltrated macrophages. While our results did confirm infiltration of peripheral macrophages (*p* < 0.05) following KA administration, there was not a statistically significant difference in the number of CD169^+^TMEM119^-^ cells between groups (**Fig. 3J, M, suppl. 3B**).

### Transcriptomic alterations in microglia upon chronic activation of Gi-signaling following *status epilepticus*

We next sought to investigate the underlying mechanisms affected by chronic Gi-Dreadd activation. To this end, RNA was extracted from hippocampal CD11b^+^ 3 days post-KA from both control and Gi-Dreadd mice (**Fig. 4A**). RNAseq revealed a total of 69 genes were differentially expressed in the Gi-Dreadd group relative to control group. Differentially expressed genes (DEGs) had an even distribution of chromosome locations (**Fig. 4B**). We also found that 43 DEGs were down-regulated within Gi-Dreadd activated mice while the remaining 26 were up-regulated (**Fig. 4C**). The heatmap shows the relative expression level of all DEGs between groups (**Fig. 4D**). Of note, the downregulated genes included several interferon-regulated genes. We also observed differential expression of Ace (Angiotensin-converting enzyme), Siglec1 (sialic acid binding Ig like lectin 1), and Mki67, a marker of proliferation. Among the fewer up-regulated, the most pertinent ones were Chrm4 (cholinergic receptor muscarinic 4), Tuba1a (Tubulin alpha 1a), Dusp8 (dual specificity phosphatase 8), and lL1α (interleukin 1 alpha) (**Fig. 4C, D**). Next, we performed Gene Ontology (GO) enrichment analysis (**Fig. 4E**). In addition to the GO analysis, we performed Gene Set Enrichment Analysis (GSEA), which determined whether the whole transcriptome was significantly different between groups (**Table 1**). We found that most significantly altered genes were involved in interferon β signaling and microglia cell-proliferation.

**Figure 4:**
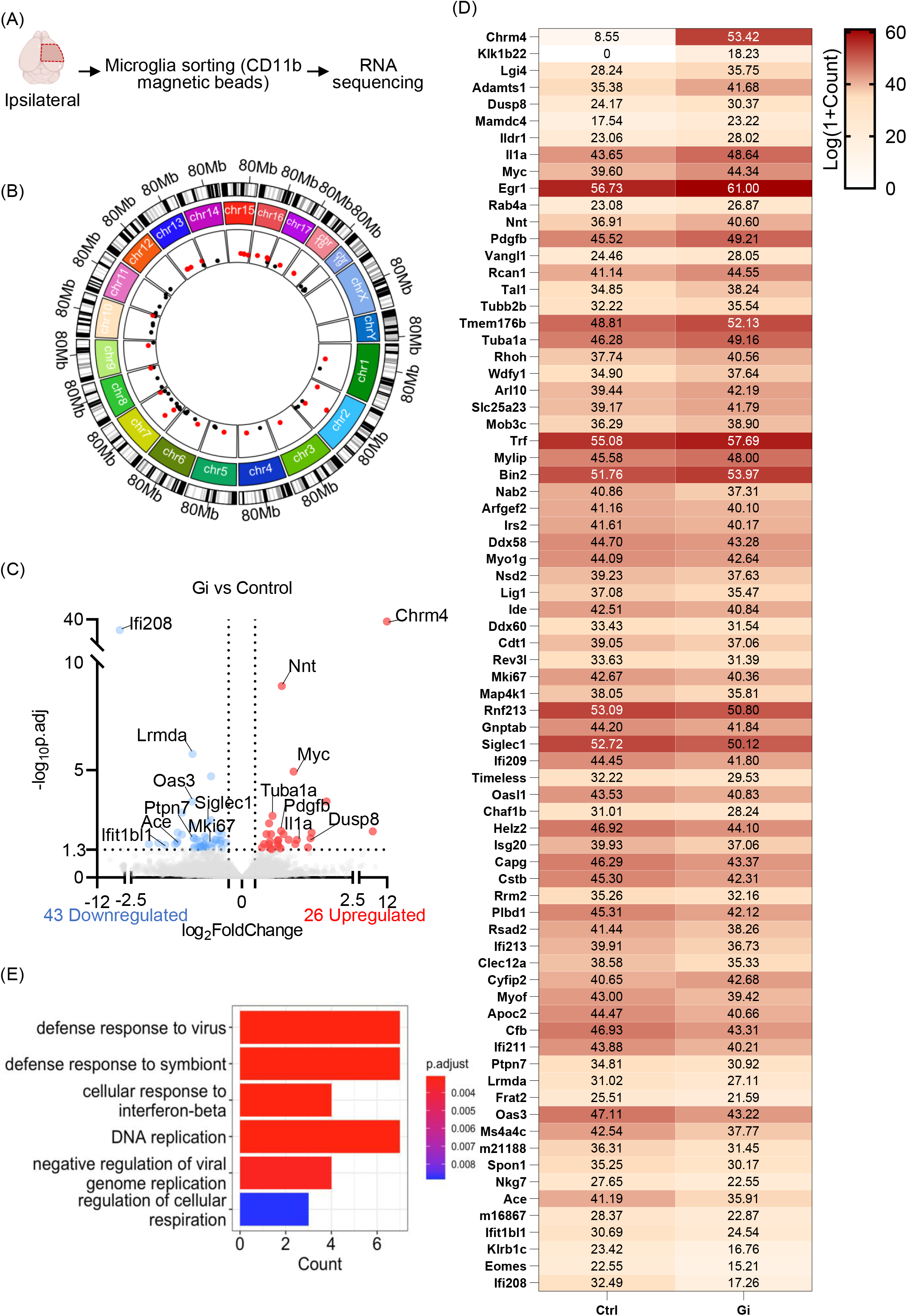
Transcriptomic alterations in microglia upon chronic Gi-signaling activation following *status epilepticus*. (A) Overview of RNAseq experiment design. (B) Visualization of differentially expressed gene (DEG) chromosomal locations. Upregulated DEGs are labeled pink and downregulated DEG are labeled black. (C) Volcano plot of DEGs. Threshold: p.adj <0.05, |log2 fold-change| > 0.5. (D) Heatmap of each DEG. Relative expression: Log(1+Count). (E) Boxplot of significant GO pathways.

**Table 1:** Top GO terms identified by GSEA.

| **GO term ID** | **Description** | **P<sub>adj.</sub>** |
| --- | --- | --- |
| GO:0035456 | response to interferon-beta | 0.000499 |
| GO:0009615 | response to virus | 0.000809 |
| GO:0051607 | defense response to virus | 0.000809 |
| GO:0140546 | defense response to symbiont | 0.000809 |
| GO:0002253 | activation of immune response | 0.002313 |
| GO:0035458 | cellular response to interferon-beta | 0.002914 |
| GO:0045089 | positive regulation of innate immune response | 0.008068 |
| GO:0002833 | positive regulation of response to biotic stimulus | 0.008068 |
| GO:0050778 | positive regulation of immune response | 0.009191 |
| GO:0002831 | regulation of response to biotic stimulus | 0.011588 |
| GO:0002218 | activation of innate immune response | 0.015124 |
| GO:0045088 | regulation of innate immune response | 0.015124 |
| GO:0060700 | regulation of ribonuclease activity | 0.019769 |
| GO:0072111 | cell proliferation involved in kidney development | 0.019769 |
| GO:0000819 | sister chromatid segregation | 0.024317 |
| GO:0009111 | vitamin catabolic process | 0.024678 |
| GO:0032069 | regulation of nuclease activity | 0.030627 |
| GO:0007059 | chromosome segregation | 0.035118 |
| GO:0006281 | DNA repair | 0.041036 |
| GO:0030550 | acetylcholine receptor inhibitor activity | 0.044216 |
| GO:1901722 | regulation of cell proliferation involved in kidney<br>development | 0.048093 |

### Prolonged activation of microglial Gi-signaling during disease progression increases neurodegeneration

Next, we determined how prolonged microglial Gi-signaling activation affected seizure associated neurodegeneration. We found a significant decrease (*p <* 0.05) in CA3 hippocampal neuronal cell number (NeuN^+^) in the Gi-Dreadd group when compared to the controls (**Fig. 5A, B**). Additionally, Nissl staining showed a significantly (*p <* 0.05) lower number of healthy, Nissl^+^ cells in the pyramidal layer of the Gi-Dreadd mice (**Fig. 5C, D**). Consistently, Fluoro-Jade C showed increased neurodegeneration in the Gi-Dreadd group (**Fig. 5E, F**). These results indicate that chronic activation of microglial Gi-Dreadd promotes the KA-induced neuronal death.

**Figure 5:**
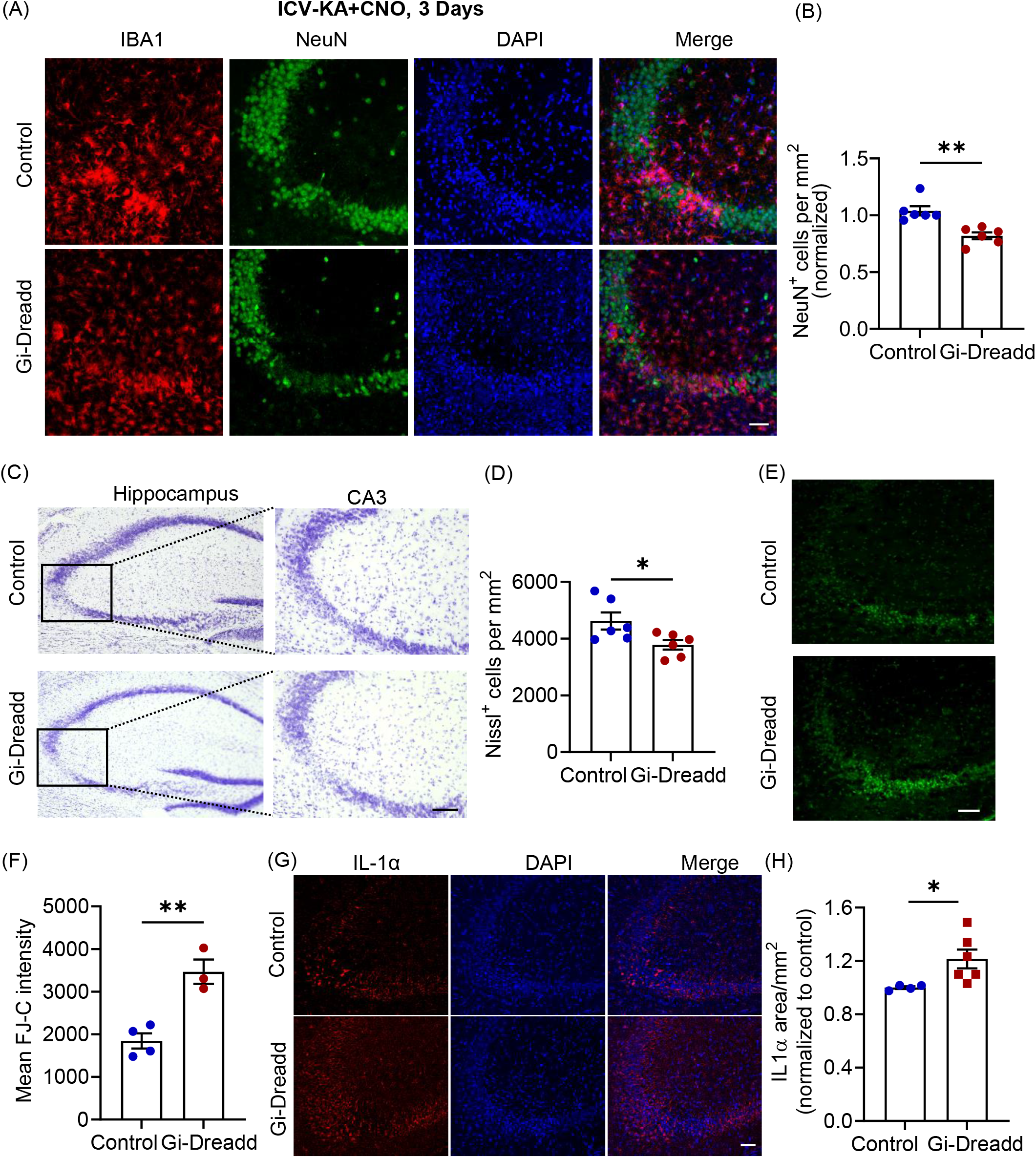
Prolonged microglial Gi-Dreadd activation causes increment in neuronal death. (A) Representative images of IBA1 (red), NeuN (green) and DAPI (blue) in the hippocampal CA3 region of Gi-Dreadd and Control mice 3 days after KA administration, scale bar, 50 µm. (B) Prolonged Gi-Dreadd activation reduced NeuN+ cells density when compared to controls. (C) Representative Nissl staining images, scale bar, 50 µm. (D) Prolonged Gi-Dreadd reduced the number of Nissl positive cells. (E) Representative images of Fluoro Jade-C (F J-C) staining, scale bar, 50 µm. (F) Prolonged Gi-Dreadd activation increased the number of FJ-C positive cells relative to controls. (G) Representative images of IL-1α (red) and DAPI (blue), scale bar, 50 µm. (H) Gi-Dreadd increased IL-1α immunoreactivity when compared with controls. Data represented as mean ± SEM, *p < 0.05, ** p < 0.01, *** p < 0.001.

Since our RNAseq results indicated an increase in the Il-1α expression within the Gi-Dreadd group, we wanted to determine if this contributed to the observed effect upon neurodegeneration. As such, we performed immunostaining for IL-1α. Our results confirmed significant upregulation (*p <* 0.05) in IL-1α expression in the Gi-Dreadd group when compared to the controls (**Fig. 5G, H**). IL-1α has been previously reported to induce activation of neurotoxic A1 type astrocytes [42], that may contribute to neuronal death. Additionally, we found that inhibiting microglia via Gi-Dreadd did not reduce astrocyte activation as determined by GFAP immunoreactivity between control and Gi-Dreadd groups (**Suppl. 4A, B, C),** suggesting that astrocytes exhibited an activated phenotype.

## Discussion

Mounting evidence describes the importance of microglia in controlling both homeostatic and pathologic brain function. In the epileptic brain, microglia adopt various phenotypes depending on the disease stage and can display both protective or deleterious functions [3, 13, 21, 43]. Using a chemogenetic approach to manipulate microglia, our results found that Gi-Dreadd activation reduced seizure severity in mice acutely. Whereas prolonged activation of Gi-Dreadd, arrested microglia in a homeostatic-like state which correlated with increased neuronal loss. Understanding the role of microglia in regulating seizures is crucial to developing potential therapeutic targets to ameliorate seizures. Our current study sheds light on microglial function and provides new avenues for future research.

### Acute activation of microglial Gi-signaling dampened seizure severity by increasing microglia-neuronal interactions

The protective role of microglia in acute seizures has been previously studied by employing genetic deletion and ablation strategies [3, 6, 12, 13, 44]. Consistently, our own results also point towards a protective role of microglia. Moreover, another study using a MgPTX mouse line showed that inhibition of Gi signaling in microglia resulted in hypersynchrony and spontaneous seizures [38]. Further, microglia can respond rapidly to acute neuronal hyperactivity [21, 45] and form purinergic junctions to monitor neuronal activity [20]. We also observed increased microglial interaction with the neurons following acute seizures in response to Gi-Dreadd activation. G protein-coupled receptor activation has been linked with cytoskeletal rearrangement [46, 47], identifying the critical function of Rho-GTPases in actin remodeling [48]. Therefore, activation of Gi-Dreadd may contribute to cytoskeletal rearrangement and increased microglial process interaction immediately following *SE*. Our results also indicated that activation of microglial Gi-signaling reduced *SE-*induced neuronal hyperexcitability. The mechanism underlying microglial neuroprotective function may be related to the production of adenosine mediated by microglia and associated suppression of neuronal activity by the adenosine receptor 1 as reported recently [19]. Our current study supports the role of Gi-induced acute microglial activation in reducing seizures.

### Chronic Gi-Dreadd stimulation inhibits microglial activation following KA induced seizures

Microglia possess both neurotoxic and neuroprotective functions [49] which greatly depend on the precise timing and context of disease [3, 50]. Curiously, when we investigated the effect of sustained Gi-Dreadd activation within microglia we found, opposite to what we observed immediately following *SE*, microglial reactivity was reduced. These results are consistent with our previous study within the context of neuropathic pain where repeated Gi-Dreadd stimulation reduced microglial activation [37]. Likewise, Binning *et al* describe the opposing effects during acute and chronic activation of Gq-Dreadd in microglial cells, showing that acute activation of Gq in microglia increased pro-inflammatory cytokine production, while this was inhibited by chronic activation [51]. Interestingly, we observed that expression of the homeostatic marker P2Y12 was elevated in the Gi-Dreadd group after *SE* when compared to controls. As such, we can conclude that chronic Gi-signaling stimulation reduces activation of microglia and locks them into a more homeostatic state. However, further study is needed.

Gi signaling could regulate downstream pathways mediated by Gi-coupled receptors like P2Y12 [52] and adenosine 1 receptor [53]. Interestingly, GO analysis highlighted the involvement of genes associated with interferon beta (IFN-β) in our study. Prior literature indicates that microglia are sensitized to IFN-β following KA induced seizures [15]. Moreover, IFN-β has been shown to regulate microglial function in multiple sclerosis [54], highlighting an anti-inflammatory role of IFN-β [55]. However, there are also clues that blocking type-I interferons may be beneficial to brain aging [56]. There is also indication of elevated IL-1α levels in the Gi-Dreadd group compared to controls. IL-1α is a pro-inflammatory cytokine [57] and has been previously reported to be an inducer for astrocytes [42]. Our results confirm the upregulated immunoreactivity of IL-1α in the Gi-Dreadd group.

We observed reduced microglia cell density following prolonged Gi-Dreadd activation. Our immunostaining results as well as RNAseq data are in accordance with these findings. Lower Mki67 levels are indicative of reduced proliferation of microglia in the Gi-Dreadd group. Additionally, increased expression of Tuba1a gene was observed in the Gi-Dreadd group as compared to the controls. Tuba1a gene encodes for the microtubule protein α-Tubulin which is essential for cytoskeleton organization and cell motility in neurons [58–60]. This supports our observation of a more ramified morphology of microglia after chronic Gi-activation.

### Inhibiting microglia during disease progression is detrimental to neuronal survival

Finally, our results suggest that arresting microglia in a homeostatic state following seizures may have a detrimental effect on neuronal health. Similarly, we found loss of TREM2 locked microglia in homeostatic states that impairs microglial protective functions in TDP-43 neurodegeneration [61]. Microglia have roles as the resident CNS immune cells, such as the removal of cell debris, and cytokine release (Sierra et al., 2010). Additionally, microglia increase expression of CD68, phagocytic function, after SE which prevents the formation of hyperexcitable aberrant neural circuits [62]. We observed reduced CD68 immunoreactivity following Gi-Dreadd activation indicating impaired removal of abnormal cells and debris. It should be noted that other cells like infiltrating monocytes [41, 63] and astrocytes also play roles in neuroinflammation and neuronal death [64]. In our study we report that there was comparable monocyte infiltration and astrocyte cell density in relation to Gi-Dreadd activation with control group. Lastly, it is worth considering that microglial activation in the acute phase may also contribute towards neuronal clearance in the Gi-Dreadd group as an initial response.

Our findings highlight a dual function for microglia Gi-signaling in epileptogenesis. Acute activation of microglia Gi-Dreadd reduces neuronal activity supporting the protective function of microglia activation in epilepsy, while, sustained activation of Gi-signaling results in microglia arrest which has detrimental outcomes. Our study strengthens the idea that microglial regulation in epilepsy is bi-phasic and may be employed as a potential therapeutic target for epilepsy prevention.

## Materials and Methods

### Animal model

Cx3cr1^CreER/WT^ (Control) and Cx3cr1^CreER/WT^: R26^LSL-hM4Di/WT^ (Gi-Dreadd) mice were used for the study. The Cx3cr1^CreER/WT^ mice were generated by crossing C57BL/6J with Cx3cr1^CreER/CreER^ (021160) mice [65]. For generating Cx3cr1^CreER/WT^: R26^LSL-hM4Di/WT^ mice, homozygous R26^LSL-hM4Di^ (026219) [66] were crossed with Cx3cr1^CreER/CreER^. Tmem119 ^CreER/WT^: R26^LSL-hM4Di/WT^ were generated by breeding Tmem119^CreER/CreER^ (031820) mice with R26^LSL-hM4Di /hM4Di^ mice. All mouse lines were obtained from the Jackson Laboratory, and then bred at Mayo Clinic. Age matched 8- to 10-week-old, male mice were used for the study in accordance with the institutional guidelines approved by animal care and use committee at Mayo Clinic.

### Gi-Dreadd expression and activation

All injection paradigms were same for both the groups. In this transgenic model Gi-Dreadd expression is controlled by tamoxifen inducible Cre expression under the microglial Cx3cr1. To induce the expression of Cre recombinase, tamoxifen (150 mg/kg) in corn oil (20 mg/mL) was injected intraperitoneally (i.p.) once per day, alternating every 2 days for a total of 4 doses. The Cx3cr1 receptor is mainly expressed in microglia but also expressed by other immune cells like macrophages [67]. Since monocytes have a fast turnover rate [65, 68], all experiments were performed after 28 days of tamoxifen injection to allow only resident microglia specific Gi-Dreadd expression. For the Tmem119 promoter induced Gi-Dreadd mouse model, experiments were performed seven days after tamoxifen injection. Clozapine-N-Oxide (CNO, Cayman Chemicals, Cat. No. 16882) was injected (i.p.) at a dose of 5 mg/kg body weight to activate Gi-Dreadds.

### KA-induced status epilepticus

Intracerebroventricular (i.c.v.) injection of kainic acid (Tocris, cat. No. 0222) 0.15 μg in 5 μL sterile PBS was administered as previously described [4, 17, 21]. Seizures were scored as per the modified Racine scale [14, 69]. Following seizure onset, CNO (5 mg/kg body weight, i.p.) was administered 20 minutes after KA injection to mice that had an onset score higher than 3, and seizure scores were recorded.

### Fluorescent immunostaining

The procedures were described previously (Mo et al., 2019). Briefly, brain samples were collected after transcardial perfusion of deeply anesthetized mice with ice cold PBS followed by 4% PFA. 30 µm slices were cut using a cryostat (Leica CM1520). Immunostaining was performed on floating sections. Sections were washed with PBS, incubated in 0.25% triton X-100 for 45min at RT. Next, sections were blocked with appropriate serum and incubated with primary antibodies as listed in table 2, overnight at 4C, followed by appropriate secondary antibody incubation at RT for an hour. After vigorous washing, sections were mounted using DAPI containing fluoromount-G mounting medium (SouthernBiotech, Birmingham, AL). Fluorescent images were collected using the LSM980 confocal microscope (Carl Zeiss Microscopy, LLC). Z-stacked images were collected. Analysis was performed with unbiased stereology.

**Table 2:** List of Antibodies

| **Name** | **Company** | **Cat#** | **Species** | **Dilution** |
| --- | --- | --- | --- | --- |
| Iba-1 | Abcam | Ab104225 | Rabbit | 1:500 |
| Iba-1 | Wako | 011-27991 | Goat | 1:300 |
| CD68 | Abcam | Ab125212 | Rabbit | 1:500 |
| TMEM119 | Abcam | 209064 | Rabbit | 1:300 |
| c-fos | CST | 2250S | Rabbit | 1:500 |
| NeuN | Abcam | Ab104225 | Rabbit | 1:500 |
| NeuN | Abcam | Ab104224 | Mouse | 1:500 |
| GFAP | CST | 3670s | Mouse | 1:500 |
| P2Y12 | Anaspec | As55043A | Rabbit | 1:400 |
| CD169 | BIO-RAD | MCA884 | Rat | 1:400 |
| Ki67 | Santa Cruz | Sc23900 | Mouse | 1:300 |
| Anti-HA-Peroxidase | Roche<br>(Sigma-Aldrich) | 23551600<br>(Clone 3F10) | Rat | 1:200 |
| GFP | Aves Labs | GFP-1010 | Chicken | 1:1000 |
| IL-1 beta | Cell signaling | 12242S | Mouse | 1:400 |
| IL-1 alpha | Abcam | Ab7632 | Rabbit | 1:200 |

### Nissl staining (cresyl violet)

Morphology of the neurons was observed by Nissl staining. The cresyl violet acetate solution is frequently used to stain Nissl substance in the cytoplasm of neurons in PFA or formalin-fixed tissue. 30 µm coronal sections from each group were air dried at room temperature overnight and processed for Nissl staining. Once rinsed with distilled water, sections were stained in 0.1% cresyl violet staining solution for 2-3 min, dehydrated using alcohol gradient of concentration of 50%, 70% and 100%, followed by clearing with xylene. Sections were then mounted using DPX mounting medium (VWR, Hatfield, PA). Light microscopy was performed using the BZ-X800 Keyence microscope (Keyence corporation of America, Itasca, IL, USA) to evaluate the morphology of the cells in CA3 region of hippocampus for both control and Gi-Dreadd groups after 3 days of i.c.v. KA induced seizures. The small-sized, dense, irregular-shaped pyknotic cells were considered as non-viable neurons.

### Fluoro jade C (FJC) staining

FJC is a poly anionic fluorescence derivative that binds with degenerating neurons with high sensitivity. The sections were stained with FJC solution (AG325, Millipore-Sigma, Burlington, MA) in 0.1% acetic acid for 30 min. The protocol followed as per user manual with slight modifications. Briefly, the sections were washed twice with distilled water and then immersed in 1% NaOH in 80% alcohol for 5 min followed by a further immersion in 70% alcohol for 2 min and then washed in distilled water for 2 min. Freshly prepared 0.06% potassium permanganate solution was added in each section for 15–30 min and gently shaken on a rotating platform. After one wash with distilled water, sections were transferred to FJC (0.0001% working solution) staining solution for 30 min in the dark, followed by subsequent washes with distilled water (1 min each). Excess water from the slides was drained off and the slides were quickly air dried on hot plate at ∼50 °C. The slides were then cleared in xylene solution and mounted in DPX mounting media (VWR, Hatfield, PA). Sections were visualized using a suitable filter system for visualizing fluorescein or FITC in the microscope (LSM 780, Carl Zeiss Microscopy, LLC). A bright green fluorescence signal indicated degenerating neurons.

### Microglia isolation and RNA extraction

Following PBS perfusion, brains from 8 weeks old mice were isolated and the ipsilateral hippocampus was extracted for microglia isolation. 3-4 hippocampi were pooled together, and 4 sets were obtained. Microglia were isolated with the magnetic CD11b cell sorting using MACS Adult Brain Dissociation kit (order no. 130-107-677, Miltenyi Biotech, Germany) as per manual instructions. Thereafter, RNA was extracted using RNeasy microkit (Qiagen, Cat. No. 74004). RNA purity and integrity was assessed, and samples were sent out for RNA sequencing to BGI company (BGI Genomics, Shenzhen, China).

### Bulk RNA sequencing and data analysis

Raw sequencing data was first filtered by the SOAPnuke software [70] for quality control purpose. Clean reads were then aligned to the *mus musculus GRCm38.p6* reference genome by the HISAT2 software [71]. Bowtie2 software was used to align clean reads to reference genes for quantification [72]. Differentially expressed genes (DEGs) between the Gi and the control group were calculated by the DESeq2 software [73]. A threshold of *p.adj <0.05* and *|log2 fold-change| > 0.5* was used to determine DEGs between conditions. Circlize software was used for the circular visualization of DEGs onto the representative genome diagram [74]. ClusterProfiler software was used for the Gene Ontology (GO) pathway enrichment analysis and the Gene Set Enrichment Analysis (GSEA)[75].

### Statistical analyses

All statistical analysis was performed using GraphPad Prism software. All results are expressed as mean ± SEM. G*Power v3.1.9.4 power analysis was performed to determine appropriate sample size. Multiple groups were compared using a two-way ANOVA. To compare results of two groups unpaired Student’s t-test were performed. P values < 0.05 are considered as statistically significant.

## Acknowledgments

This work was supported by the following grants from the National Institutes of Health: R01NS088627 (L.-J.W.) and R35NS132326 (L.-J.W.).

## Author contributions

A.D. and L.J.W. designed the study and wrote the manuscript. A.D. performed the experiments and collected data. D.B.B assisted with experimental design. J. Z. and S.Z. assisted with RNA sequencing experiments. L.W. performed data mining and RNAseq data analysis. K.H. helped with data analysis. M.H.Y., A.B. and D.T. assisted with manuscript revision.

## Declaration of interests

The authors declare no competing interests.

---

**Suppl. Figure 1:**
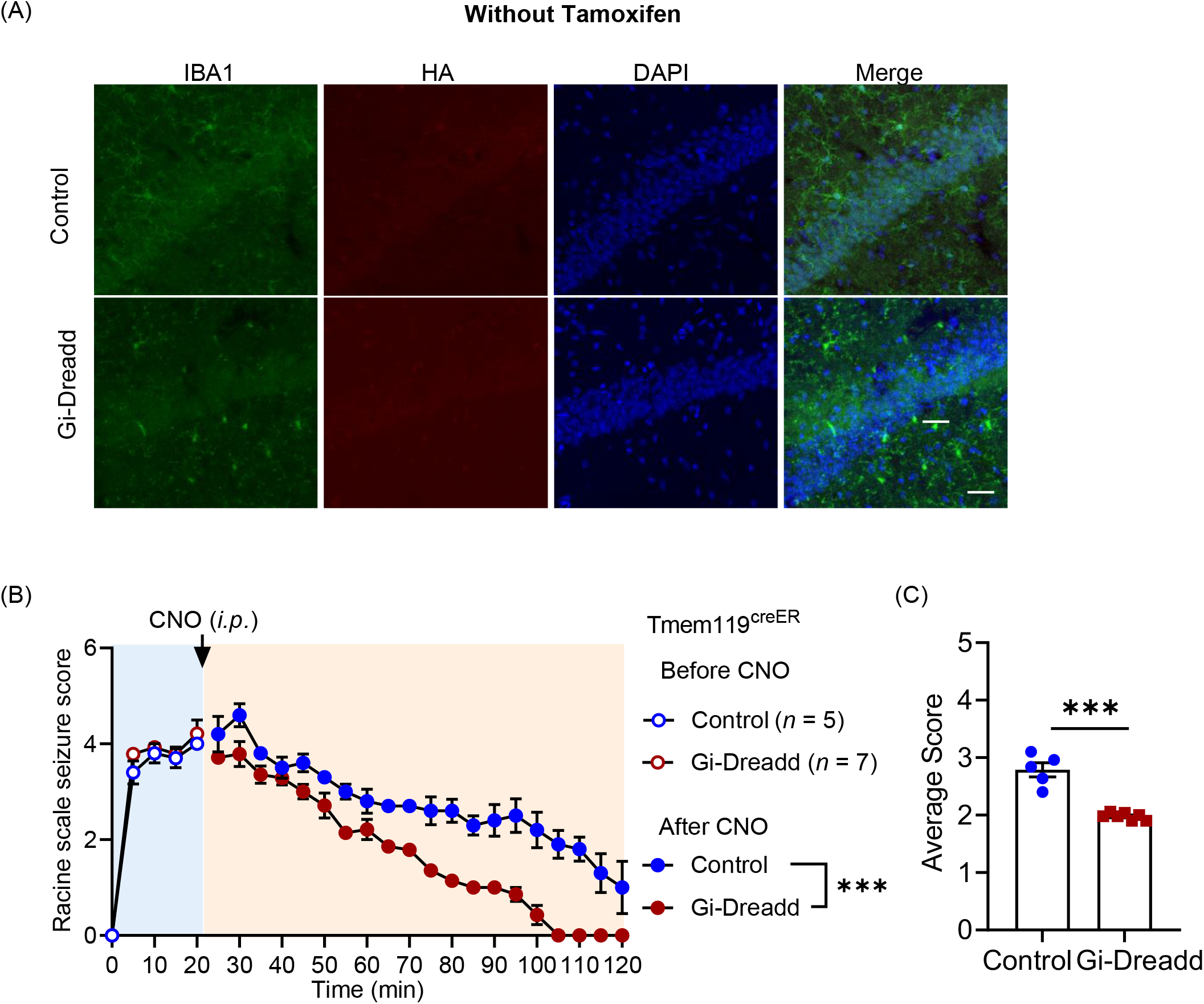
Chemogenetic modulation of microglia reduces seizures confirmed by Tmem119Cre induced Gi-expressing mice. (A) Representative immunostaining images depicting no HA expression in Gi-Dreadd mice when tamoxifen is not administered confirming the efficacy of cre dependent Gi-Dreadd expression. Scale bar 30μm. (B) Seizure profile for mice that obtained a score of 3 and above in the first 20 min after KA administration. Results indicate that Gi-Dreadd activation by CNO reduced the severity of seizures. Analyzed by two-way ANOVA. (C) Graph representing average seizure score showing significantly reduced score in the Gi-Dreadd group. Analyzed by t-test. Data represented as mean ± SEM, *p < 0.05, ** p < 0.01, *** p < 0.001.

**Suppl. Figure 2:**
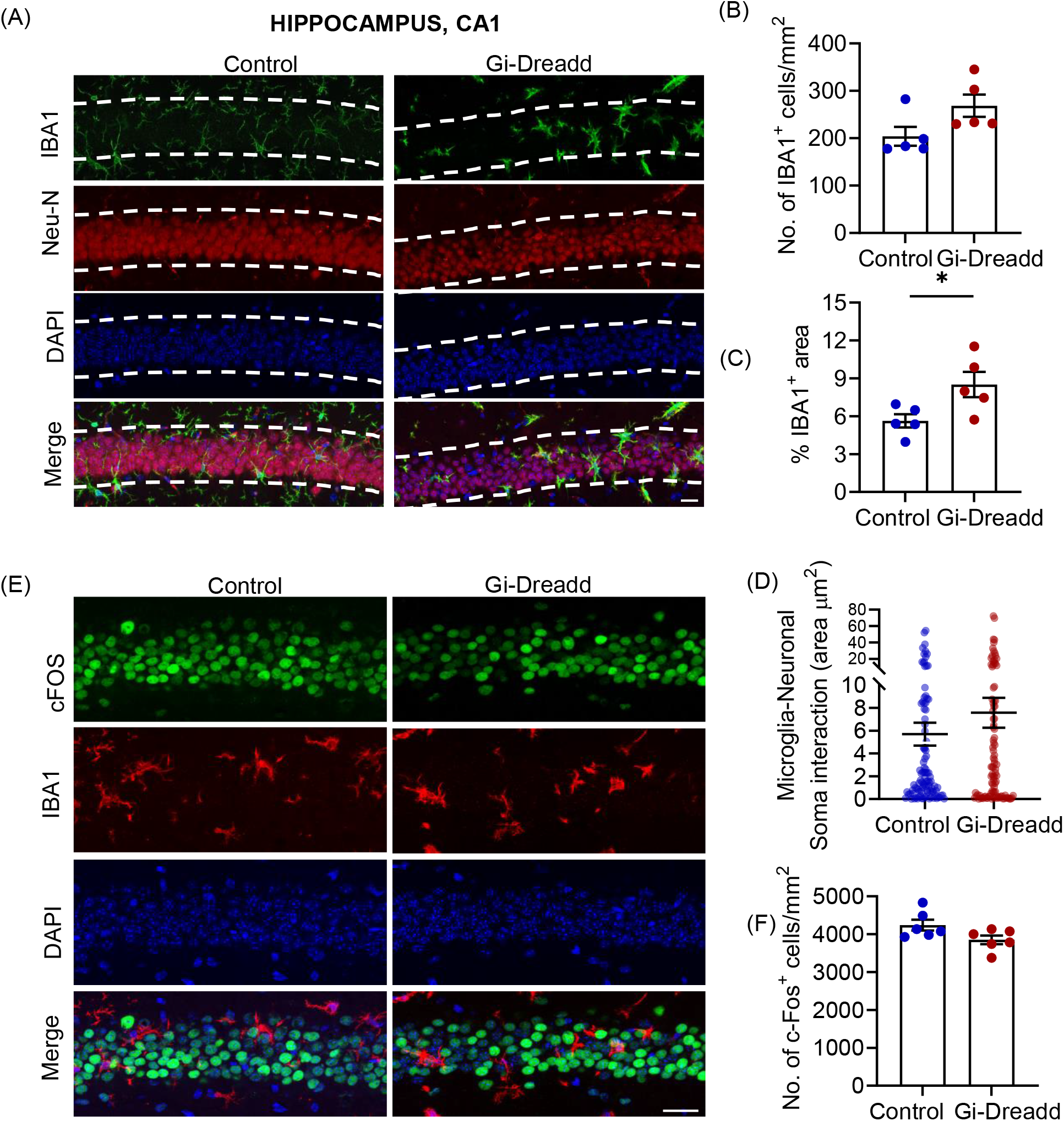
Effect of activation of microglial Gi-Dreadd by CNO following *status epilepticus* in CA1 region of hippocampus. (A) Representative immunostaining images of IBA1 (green), NeuN (red). and DAPI (blue) in CA1 region of hippocampus of Gi-Dreadd and Control mice. Scale bar, 50 µm. (B) Graph represents no difference in number of IBA1 positive cells, however, (C) IBA1 immunoreactive area in the Gi-Dreadd group is elevated when compared to Control. (D) Analyzed results show no difference in microglia interaction with Neuronal soma between the two groups in CA1 following acute seizures. (E) Images for c-FOS (green), IBA1 (red), and DAPI (blue), scale bar, 30 µm. (F) Summarized results show no difference in CA1 region. Data represented as mean ± SEM, *p < 0.05.

**Suppl. Figure 3:**
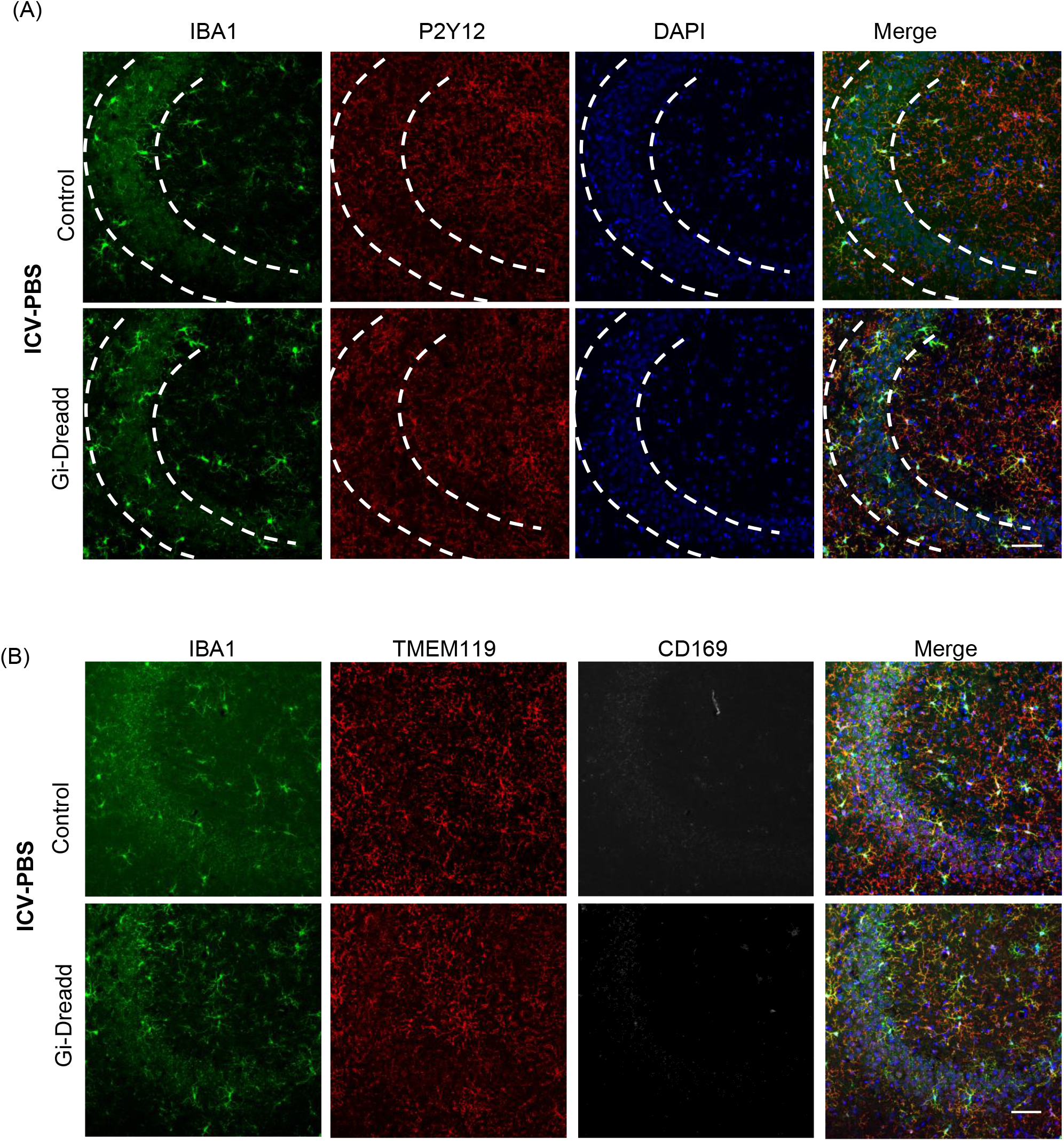
Effect of activation of microglial Gi-Dreadd by CNO following *status epilepticus* in CA1 region of hippocampus. (A) Representative images of IBA1 (green), P2Y12 (red), and DAPI (blue) in CA3 region of hippocampus of i.c.v.-PBS+CNO injected Gi-Dreadd and Control mice. (B) Images for IBA1 (green), Tmem119 (red). and DAPI (blue) from i.c.v.-PBS+CNO injected Gi-Dreadd and Control mice (scale bar, 50 µm).

**Suppl. Figure 4:**
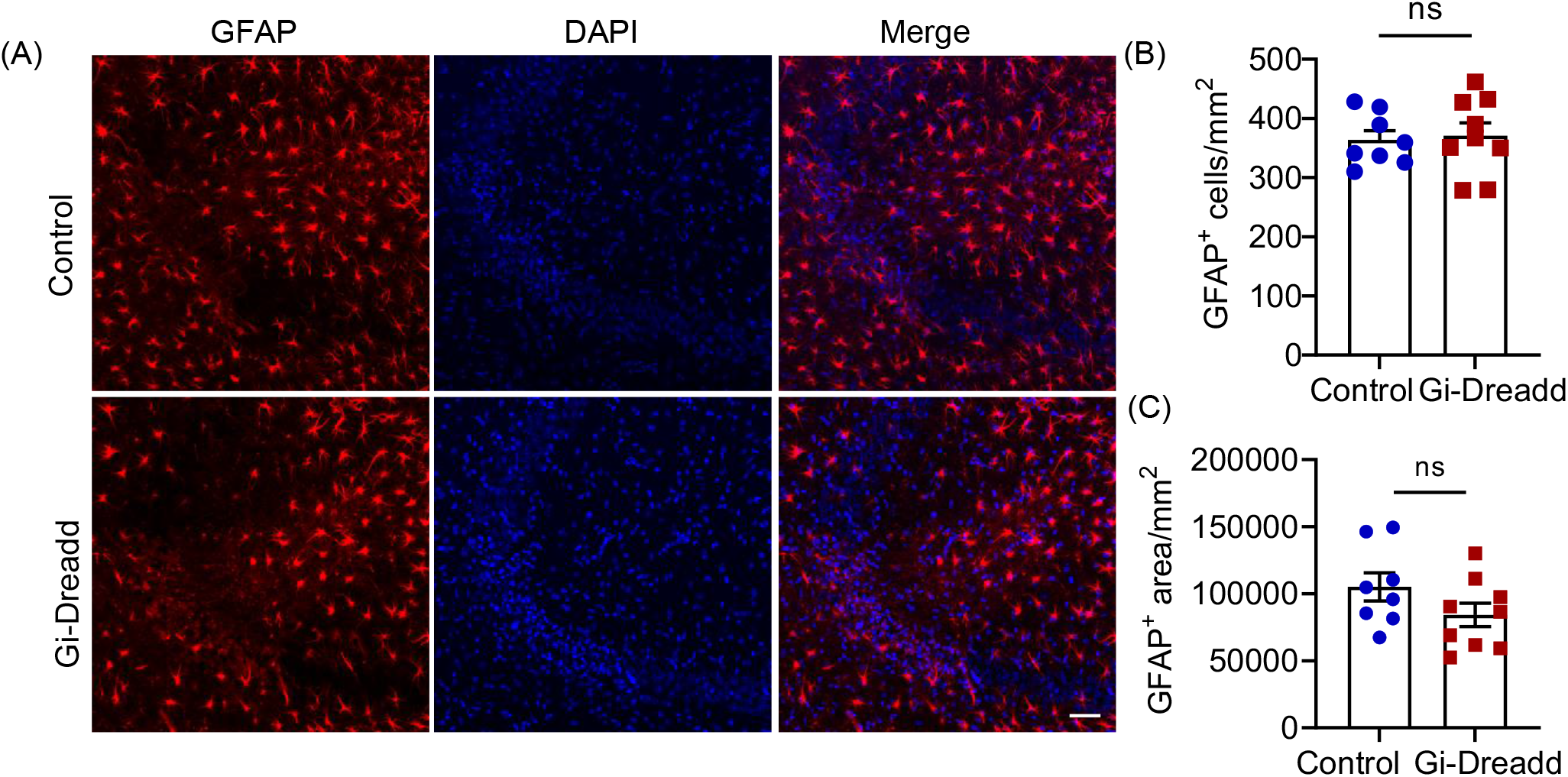
Immunohostochemical analysis of astrocyte activation following microglial Gi-Dreadd activation by CNO. (A) Representative immunostaining images of GFAP (red) and DAPI (blue) in CA3 region of hippocampus of i.c.v.-KA+CNO injected Gi-Dreadd and Control mice. (B) Graph representing GFAP+ cell number in response to seizures. (C) Graph depicting GFAP immunoreactive area (scale bar, 50 µm).

## Notes

### Competing Interest Statement

The authors have declared no competing interest.

